# ZygosityPredictor

**DOI:** 10.1101/2023.03.09.531877

**Authors:** Marco Rheinnecker, Martina Fröhlich, Marc Rübsam, Nagarajan Paramasivam, Christoph E. Heilig, Stefan Fröhling, Richard F. Schlenk, Barbara Hutter, Daniel Hübschmann

**Affiliations:** Computational Oncology Group, Molecular Precision Oncology Program, National Center for Tumor Diseases (NCT) Heidelberg and German Cancer Research Center (DKFZ); German Cancer Consortium (DKTK) Heidelberg; Division of Translational Medical Oncology, NCT Heidelberg and DKFZ; Department of Medical Oncology, NCT Heidelberg, Heidelberg University Hospital; Department of Hematology, Oncology and Rheumatology, Heidelberg University Hospital; NCT Trial Center, NCT Heidelberg, Heidelberg University Hospital and DKFZ; Pattern Recognition and Digital Medicine Group, Heidelberg Institute for Stem Cell Technology and Experimental Medicine (HI-STEM)

## Abstract

**Summary:** ZygosityPredictor provides functionality to evaluate how many copies of a gene are affected by mutations in next generation sequencing data. In cancer samples, the tool processes both somatic and germline mutations. In particular, ZygosityPredictor computes the number of affected copies for single nucleotide variants and small insertions and deletions (Indels). In addition, the tool integrates information at gene level via phasing of several variants and subsequent logic to derive how strongly a gene is affected by mutations and provides a measure of confidence. This information is of particular interest in precision oncology, e.g. when assessing whether unmutated copies of tumor-suppressor genes remain.

**Availability and implementation:** ZygosityPredictor was implemented as an R-package and is available via Bioconductor at https://bioconductor.org/packages/ZygosityPredictor. Detailed documentation is provided in the vignette including application to an example genome.

## 1 Introduction

Variant calling is the process of detecting mutations in data obtained from sequencing nucleic acids in a biological sample [4]. When working with short-read next generation sequencing (NGS), variant calling is performed down-stream of alignment of the reads to the reference genome. Variant calling can be performed for both germline variants, which are present in all cells of an organism, and for somatic variants, which occurred during the life span of the organism and which are present only in clonally expanded cells like, e.g., in a neoplasm. Variant calling can be performed for different mutation types, including single nucleotide polymorphisms (SNPs)/single nucleotide variants (SNVs), small insertions and deletions (Indels), copy number variations (CNVs), somatic copy number aberrations (sCNAs) and structural variants (SVs) [4, 8]. In the following, we designate SNVs and Indels together as small variants. Detection of small variants is almost always based on criteria of read counts or allele frequency. When analyzing cancer samples, read counts are modulated by the copy number state of the genomic segment in which the small variant is located in, as well as many other factors like tumor cell content, i.e., the purity of the sample. However, many strategies for small variant calling don’t make use of the local copy number information. In a precision oncology setting, this may have strong consequences on downstream processing, including identification of driver gene aberrations and targetable lesions, and on clinical decision making [2]. For correct interpretation of the functional state of a mutated gene, it is important to determine how many copies of that gene are affected by a mutation (*affected copies, ac*). In particular, it is of high relevance if all copies are affected (bi-allelic mutations in the diploid case) or if unmutated copies remain (mono-allelic mutations in the diploid case). Furthermore, if a gene is affected by more than one mutation, e.g. in the case of compound heterozygosity, it is important to phase the mutations, to determine whether they affect the same allele or different ones. Here, we present ZygosityPredictor, an R/Bioconductor package with functionality for the computation of the number of affected copies (downstream of small variant and CNV / sCNA calling) for both germline and somatic mutations as well as for phasing. Phasing is performed on DNA sequencing data and, if available, on RNA-seq data, making use of both read-level phasing information and haploblocks. These data layers are integrated in order to provide information whether all copies of a gene are affected or whether unmutated copies remain. An important modulating factor for all allele-frequency-derived measures is the presence of subclones. In bulk-sequencing, accounting for subclonality is difficult. In this work, we assume that the investigated tumor is clonal. More precisely, we assume that the cancer cell fraction (CCF), i.e., the fraction of the tumor cells that carry the variant, is 1 [9]. This is a clear limitation of our software, and violations of this assumption may occur in real data. Of note, our software assembles a versatile toolbox, not all of which is novel. The novelty is primarily in determining the mutation status of genes.

**Figure 1.**
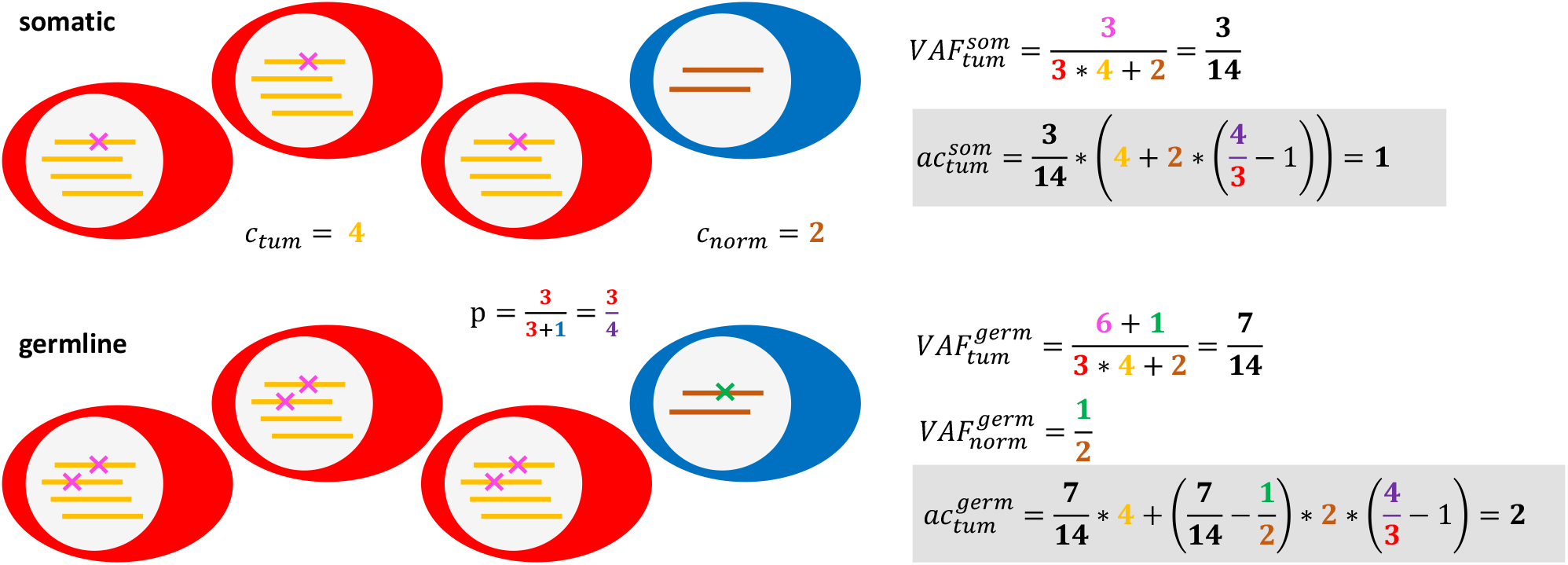
Schematic for determination of the number of affected copies for a somatic or germline mutation in a tumor sample. Tumor cells are depicted in red, admixed healthy cells are depicted in blue. The relevant parameters are color-coded. Calculation of affected copies are depicted in grey boxes.

## 2 Methods and Results

ZygosityPredictor is to be executed after calling of small variants (germline and somatic) [3, 1] and sCNAs [5]. By integrating information at gene level, ZygosityPredictor computes if all copies are affected (bi-allelic mutations in the diploid case) or if unmutated copies remain (mono-allelic mutations in the diploid case). To make the problem identifiable, we make some restrictive assumptions: The provided variants and copy number profiles must originate from a clonal tumor, i.e., the fraction of the tumor cells that carry the variant, is 1. The interpretability of the results increases if the input variants are filtered to the functionally relevant ones. ZygosityPredictor generally operates in two steps: calculation of affected copies for each variant and subsequent assessment of a gene by integration of information on all variants in this gene. Ultimately, a gene status can be assigned to each gene which is one of: *all copies affected, wt copies left* or *undefined*. Of note, the functionality of ZygosityPredictor is also applicable in cases with ploidy higher than 2, which is often the case in tumor samples.

### 2.1 Calculation of affected copies

To determine the contribution of each variant to the affection of a gene, it is necessary to estimate how many copies are affected by the mutation. For small variants, the number of affected copies is computed according to formulae (1) and (2) below. Derivations for these formulae are provided in the supplement. Let *ac*_*tum*_ define the number of affected copies in the tumor (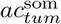 for somatic variants and 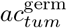 for germline variants), *c*_*tum*_ the copy number in the tumor of the genomic segment the variant is located in and 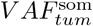 the allele-frequency of the somatic variant in the tumor sample. Let *c*_*norm*_ define the expected copy number in normal tissue of the segment the variant is located in. This will be 2 for genes located on autosomes and for genes on the X chromosome in female samples, and 1 for genes on the X and Y chromosomes in male samples (without the pseudoautosomal regions). With the definition of *p* as the purity of the tumor, we can deduce that for a somatic variant in the tumor sample, the number of affected copies is:

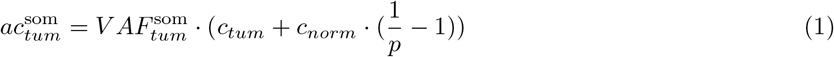

Using 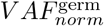, the allele-frequency of the germline variant in the normal control, and 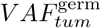, the allele-frequency of the germline variant in the tumor sample, we can compute:

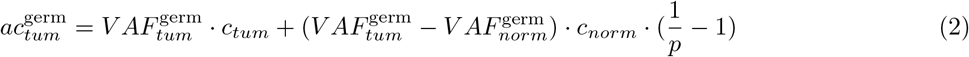

In case 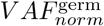 is unknown, it can be assumed to be 0.5 for heterozygous variants in genes located on autosomes or on the X chromosome in female samples, and 1 for homozygous variants and variants in genes on the X and Y chromosomes in male samples (without the pseudoautosomal regions), for a normal chromosome set.

### 2.2 Definition of gene status

To translate the number of affected copies into a gene status, the following logic and arithmetics are applied. Due to the diploid nature of the genome, mutations cannot propagate from one allele to the other. Therefore, beside the unlikely case of two identical events on both alleles, a single small variant can only affect all copies of a gene if the other allele was lost during tumor development. Therefore, the decisive criterion for a single variant to affect all copies is the detection of loss of heterozygosity (LOH). Single variants in genes without LOH will therefore always get *wt copies left*. These assumptions are motivated by the fact that pipelines used for detection of sCNAs already apply sophisticated logic for the assignment of the copy number and/or LOH state of every segment. If LOH is detected at the position of a somatic variant, the left wt copies are calculated *tum* via 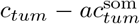 and must be below 0.5 to lead to the gene status of *all copies affected*. For germline variants in the context of LOH the decision is easier as they are to be expected on every copy originating from the same allele. Therefore it suffices to check which allele was lost in the tumor. If the allele frequency of the variant is below 0.5, we assume that the corresponding allele was lost in the tumor, if it exceeds 0.5, all copies are affected. Furthermore, in ZygosityPredictor, affection of a gene by a homozygous deletion always leads to *all copies affected*.

If up to this point none of the single variants leads to the affection of all copies of the gene, compound heterozygosity is checked. To qualify for this, several variants must be present in the same gene, which might then contribute to a constellation in which all copies of the gene are affected. To accurately assess the interplay of all variants in a gene, phasing is used (see section 2.3 below). Depending on the phasing result, the affected copies of all contributing single variants are aggregated into a quantity termed *integrated affected copies* (*iac*). Ultimately, if *c*_*tum*_ *™ iac* leaves less than 0.5 wt copies, the status *all copies affected* is attributed to the gene. Since the affected copies for each variant are already known, *iac* can be calculated for all pairwise combinations of variants. If for none of those pairs *iac* falls below 0.5 wt copies, it can be concluded that there are still wt copies left. For the purpose of confidence estimation (see section 2.4 below), such cases are flagged.

### 2.3 Phasing of variants

To reconstruct the interplay between all variants in a gene, pairwise combinations of all variants are assessed. Possible constellations for a pair of variants are either *same*, if the two variants are located on the same copy or *diff* if they are on different copies. The primary approach is read-level phasing. Here, sequencing reads or read-pairs from DNA-sequencing and, if available, RNA-sequencing, which map to the positions of both variants are checked for the presence of these variants. If there are reads that simultaneously carry both variants, it can be concluded that they originate from the same copy (constellation: *same*; *iac* = *max*(*ac*_1_, *ac*_2_)). If no reads carrying both variants are present, the variants most likely originate from different copies (constellation: *diff* ; *iac* = *ac*_1_ + *ac*_2_). As shown in (Figure 2 A), every read overlapping both positions is classified into one of four read-categories: *both*, indicating the presence of both variants, *none* indicating none of them and *mut1* or *mut2* for only one of the two variants. For the two possible constellations *same* and *diff*, different counts of the read categories are expected. For *same*, only reads carrying both or none of the variants are expected while the ones with only a single variant should not occur (*same*: *n*_*both*_ *≥* 1, *n*_*mut*1_ = 0, *n*_*mut*2_ = 0). In the *diff* case, expectation is inverted (*diff* : *n*_*both*_ = 0, *n*_*mut*1_ *≥* 1, *n*_*mut*2_ *≥* 1). For ease of nomenclature, we define successful read-level phasing of two variants as *direct phasing*. An example can be found in Figure 2 B for the variants m1 and m2 via the blue and pink read-pairs. Naturally, chances to successfully perform direct phasing decrease with increasing distance between the variants due to limited read length. To a certain extent and if available, RNA reads might help to overcome this limitation, but are themselves limited to expressed regions and may be subject to alternative splicing. If direct phasing does not succeed, indirect approaches via other mutational events like SNPs in close proximity or variants in addition to the investigated pair are initialized. Indirect phasing requires a read-level phasing of both mutations of interest to at least one such event. If a connection of both variants can be found via other mutational events in between, the constellation for the pair of interest can be concluded from the constellations with these events. In Figure 2 B, such a connection is present for m1 and m4; m1 can be phased to m3 via the RNA read, which can be phased to s4 and finally to m4 via the yellow and grey read-pairs.

**Figure 2.**
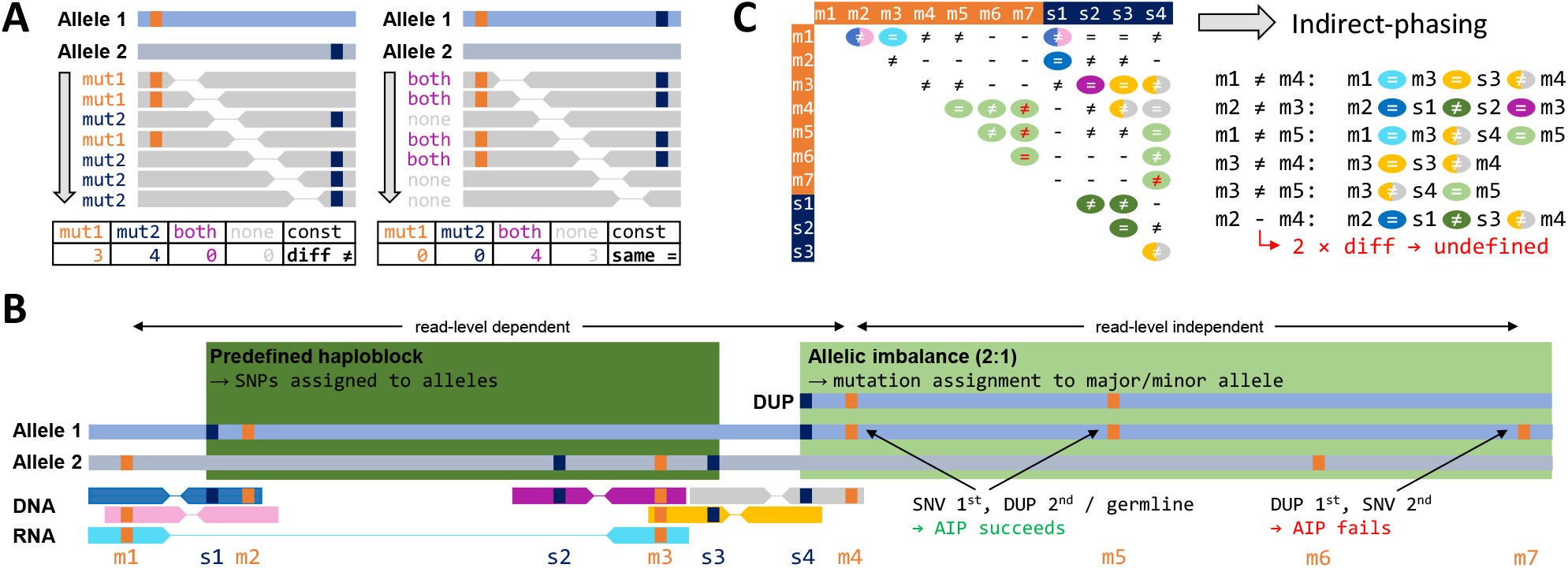
Phasing. A) Schematic for read-level phasing in a diploid genome with two variants in a region of interest. On the left, the variants are located on different alleles; on the right, variants are located on the same allele and therefore leave one wt allele intact. Below, repercussions on NGS reads/read-pairs are shown. B) Example of a genomic region with 7 mutations/variants to be phased. Variants are depicted in orange and SNPs (germline variation including polymorphisms) in blue. Abbreviation: AIP = allelic imbalance phasing. C) Phasing-matrix of the same example as in B with detailed indirect reasoning for all pairwise variant combinations that cannot be determined by direct phasing. Colors of phased combinations correspond to read-pair colors in B.

Another way for bridging long distances between variants is the use of haploblocks as computed by upstream tools, e.g. sCNA calling. We define a haploblock as a segmented genomic region in which SNPs can be assigned to one allele. If both variants can be phased to SNPs of the same haploblock, their constellation can be determined. This case is depicted for m2 and m3 in Figure 2 B, which are phased to s1 and s2, respectively, located on different alleles of the same haploblock.

If read-level phasing approaches fail, allelic imbalance phasing (AIP) can be used (Figure 2 B, m4, m5, m6, m7). Segments of allelic imbalance are characterized by deviating copy number between the two alleles. Genotypes for the variants in such segments are defined via the concept of genotype likelihood (cf. supplement) primarily based on allele-frequencies [6]. In contrast to germline variants, allele-frequencies of somatic mutations strongly depend on the sequence of mutational events during tumor development, in particular the sequence of occurence of sCNAs and small variants. If in a given segment a small variant has occurred after an sCNA, the allele-frequency of the small variant cannot be used to accurately predict allele specific genotypes (variant m7). In light of these uncertainties, we consider this approach as generally low confidence and disable it by default - it is, however, available to the expert user and can be re-activated.

Figure 2 C displays in a matrix the constellations that could be defined via all phasing approaches, using the same color code as in B. Based on these, other combinations can be imputed indirectly. However, in cases in which the inference includes two or more *diff* -constellations, we cannot confidently exclude the presence of additional copies and classify the constellation as *undefined*, as in the case of m2 and m4.

### 2.4 Confidence estimation

To quantify the reliability of the assignment of a gene’s status, ZygosityPredictor provides an aggregated confidence measure. The proposed arithmetics work as follows: a gene whose status is directly determined by a single variant or assessed by logic, i.e., either by using an LOH call or for a flagged variant (Section 2.2), gets the highest possible confidence. Lower confidence levels are attributed to genes that were assessed via phasing. For those, we start with attributing a confidence value (*c*_*rc*_) to each read classification (*both, mut1, mut2*). Let *p*_*bc*_ be the error probability of a basecall and *p*_*mq*_ the mapping quality of the read. The confidence for a read classification can then be determined according to Formula 3

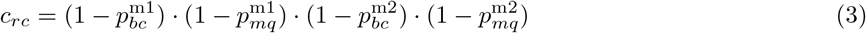

To discriminate between *same* and *diff* constellations, we use the *χ*^2^-distribution [7] to compare the obtained reads with expected ones (*same*: *n*_*both*_ *≥* 1, *n*_*mut*1_ = 0, *n*_*mut*2_ = 0 and *diff* : *n*_*both*_ = 0, *n*_*mut*1_ *≥* 1, *n*_*mut*2_ *≥* 1). We weigh each read by its *c*_*rc*_. Low quality reads or low confidence basecalls will therefore contribute less. Of note, this is not a *χ*^2^ test in the strict sense of the term; instead we present an empirical confidence measure inspired by the statistical test. Finally, for a given pair of variants, the confidence measures *c*_*indirect*_ (corresponding to 1*™p* of all these tests) of all phasing combinations used to obtain the constellation of the variant pair of interest, including those with and between SNPs (if indirect-phasing was used), must be aggregated to one confidence measure for the variant pair (*c*_*comb*_). This is done by taking the product of all *c*_*indirect*_.

For the ultimate confidence aggregation step from the pairwise variant constellations to the gene status, different considerations apply depending on the gene status. Mathematically, this aggregation is simply the product of *c*_*comb*_ of all combinations that were necessary to define the gene status. For the status *all copies affected*, a single combination that affects all copies is sufficient, while for the status *wt copies left*, every combination is required to ensure that none of them affects all copies. The final confidence estimation for the gene status is a numeric value between 0 and 1, where 1 is the maximum possible confidence. Of note, when using haploblock phasing, ZygosityPredictor is agnostic to which algorithmic strategy may have been employed for computing haploblocks and therefore instead of assigning a numeric confidence value to this category of information, a placeholder is being used (also as a factor in the composite formulae) which may be filled by the user, e.g., by inheriting from the confidence of the algorithm which has performed the haploblock computation. In case AIP has been used, confidence is estimated with the help of genotype likelihoods (cf. supplement).

## 3 Conclusion

We developed the R/Bioconductor package ZygosityPredictor which downstream of variant calling computes the number of affected copies for every gene, taking into consideration both somatic and germline variants. In particular, it computes the number of affected copies for small variants (SNVs and Indels). Then, in a second step the tool performs haplotype phasing based on DNA and, if available also RNA sequencing data, and assigns a confidence value to the phasing results. ZygosityPredictor then finally assigns categories of whether functional copies of a gene remain, which is of particular use, e.g. in precision oncology for the assessment of tumor-suppressor genes.

## 4 Funding

This work was supported by the NCT Molecular Precision Oncology Program. MRu was supported by the DKTK Joint Funding Program. MRh was funded by allowances for scientific assistant from the graduate school HIDSS4health.

## Notes

### Competing Interest Statement

The authors have declared no competing interest.

### Summary of Updates

The tool was updated with improved phasing approaches as well as confidence estimation. The manuscript was updated accordingly.

