## Supplementary information for "ZygosityPredictor"

### Supplementary Information for the manuscript "ZygosityPredictor"

#### 1 Calculation of affected copies

Let $ac_{tum}$ define the number of affected copies in the tumor ($ac_{tum}^{\text{som}}$ for somatic variants and $ac_{tum}^{\text{germ}}$ for germline variants), $c_{tum}$ the copy number in the tumor of the genomic segment the variant is located in and $VAF_{tum}^{\text{som}}$ the allele-frequency of the somatic variant in the tumor sample. Let $n_{tum}$ and $n_{norm}$ designate the number of tumor and normal cells in the sample, respectively. Then, in case of a 100 % pure tumor, the allele frequency of a somatic variant is defined as:

$$VAF_{tum}^{\text{som}} = \frac{ac_{tum}^{\text{som}} \cdot n_{tum}}{c_{tum} \cdot n_{tum}} = \frac{ac_{tum}^{\text{som}}}{c_{tum}} \tag{1}$$

In case the purity is not 100 %, the formula needs to be extended. Let $c_{norm}$ define the expected copy number in normal tissue of the segment the variant is located in. This will be 2 for most genes located on autosomes and for genes on the X chromosome in female samples, and 1 for genes on the X and Y chromosomes in male samples (without the pseudoautosomal regions). Formula 1 can then be extended as follows:

$$VAF_{tum}^{\text{som}} = \frac{ac_{tum}^{\text{som}} \cdot n_{tum}}{c_{tum} \cdot n_{tum} + c_{norm} \cdot n_{norm}} \tag{2}$$

The purity of a tumor sample is defined by:

$$p = \frac{n_{tum}}{n_{tum} + n_{norm}} \tag{3}$$

From this, we can deduce $n_{norm} = n_{tum} \cdot \frac{1-p}{p}$ and:

$$VAF_{tum}^{\text{som}} = \frac{ac_{tum}^{\text{som}}}{c_{tum} + c_{norm} \cdot (\frac{1}{p} - 1)} \tag{4}$$

and thus

$$ac_{tum}^{\text{som}} = VAF_{tum}^{\text{som}} \cdot (c_{tum} + c_{norm} \cdot (\frac{1}{p} - 1)) \tag{5}$$

In the tumor sample, the allele frequency of germline variants can be defined as:

$$VAF_{tum}^{\text{germ}} = \frac{ac_{tum}^{\text{germ}} \cdot n_{tum} + ac_{norm}^{\text{germ}} \cdot n_{norm}}{c_{tum} \cdot n_{tum} + c_{norm} \cdot n_{norm}} \tag{6}$$

Of note, $ac_{tum}^{\text{germ}}$ and $ac_{norm}^{\text{germ}}$ define numbers of affected copies for the germline variant in tumor and normal tissue, respectively. Using $VAF_{norm}^{\text{germ}}$, the allele-frequency of the germline variant in the normal control, we can substitute $ac_{norm}^{\text{germ}} = VAF_{norm}^{\text{germ}} \cdot c_{norm}$ and obtain:

$$VAF_{tum}^{\text{germ}} = \frac{ac_{tum}^{\text{germ}} \cdot n_{tum} + VAF_{norm}^{\text{germ}} \cdot c_{norm} \cdot n_{norm}}{c_{tum} \cdot n_{tum} + c_{norm} \cdot n_{norm}} \tag{7}$$

When applying the same arithmetics as above, we get:

$$VAF_{tum}^{\text{germ}} = \frac{ac_{tum}^{\text{germ}} + VAF_{norm}^{\text{germ}} \cdot c_{norm} \cdot (\frac{1}{p} - 1)}{c_{tum} + c_{norm} \cdot (\frac{1}{p} - 1)} \tag{8}$$

and thus

$$ac_{tum}^{\mathrm{germ}} = VAF_{tum}^{\mathrm{germ}} \cdot c_{tum} + (VAF_{tum}^{\mathrm{germ}} - VAF_{norm}^{\mathrm{germ}}) \cdot c_{norm} \cdot (\frac{1}{p} - 1) \quad (9)$$

In case $VAF_{norm}^{\mathrm{germ}}$ is unknown, it can be assumed to be 0.5 for heterozygous variants in most genes located on autosomes or on the X chromosome in female samples, and 1 for homozygous variants and variants in genes on the X and Y chromosomes in male samples (without the pseudoautosomal regions), for a normal chromosome set.

### 2 Allelic Imbalance Phasing (AIP)

If read-level dependent phasing approaches fail, segments of allelic imbalance can be used to determine a constellation of two variants. In such segments, sCNAs have taken place during tumor development and the number of copies of one allele differs from the number of copies of the other allele. In NGS this is reflected by deviating allele frequencies depending on which allele a variant is located on. If, for example, a genomic segment was called with allelic imbalance of 1:2, i.e., one allele is present once while the other allele was duplicated, we expect variants in this segment to have either a low or high allele-frequency, depending on whether they are on the major or the minor allele. In this work, we use genotype likelihoods to determine to which of the two cases a given variant belongs. By using formula 10 from [1], the likelihood of a variant to be located on the allele with the respective genotype can be defined.

Let $a$ denote the number of reads supporting the alternative allele at a position of a variant and $r$ the number of reads supporting the reference allele. The genotype is denoted by $g$, i.e., the number of copies of the allele which is currently checked. $\epsilon_j$ is the error probability in read $j$, i.e. $\epsilon_j$ is defined by $(1 - p_{mq,j}) * (1 - p_{bq,j}) = 1 - \epsilon_j$ where $p_{mq,j}$ is the mapping quality and $p_{bq,j}$ is the base quality. Then, the genotype likelihood of the genotype $g$ is defined as:

$$\mathcal{L}(g) = \frac{1}{(c_{tum})^{a+r}} \Pi_{j=1}^{r}[(c_{tum} - g)\epsilon_j + g(1 - \epsilon_j)]\Pi_{j=r+1}^{a+r}[(c_{tum} - g)(1 - \epsilon_j) + g\epsilon_j] \quad (10)$$

Formula 10 can now be used to determine the genotype with the highest likelihood for a given position which may contain a mutation. To determine the constellation of two variants $m1$ and $m2$ to each other, four genotype likelihoods are calculated. $gt1$ and $gt2$ denote the two possible genotypes of the segment of allelic imbalance. Likelihood for the two possible constellations can be calculated according to:

$$L_{diff} = \mathcal{L}_{m1}(gt1) * \mathcal{L}_{m2}(gt2) + \mathcal{L}_{m1}(gt2) * \mathcal{L}_{m2}(gt1) \quad (11)$$

$$L_{same} = \mathcal{L}_{m1}(gt1) * \mathcal{L}_{m2}(gt1) + \mathcal{L}_{m1}(gt2) * \mathcal{L}_{m2}(gt2) \quad (12)$$

The constellation of higher likelihood is selected. For reasons of numeric stability, we use the log-likelihood ratio:

$$LLR = log(\frac{L_{same}}{L_{diff}}) \quad (13)$$

The absolute value of the log-likelihood ratio is used as confidence measure for AIP cases. Of note, the confidence measures of AIP and read-level phasing cannot be directly compared - which, in our opinion, should not be done anyways. Both information are annotated in separate columns of the output of ZygosityPredictor.

As we assume that our samples are not fully pure, i.e the tumor cell content is lower than 100 %, we need to slightly adjust the formula for the computation of genotype likelihood.

#### 2.1 Somatic variants

For somatic variants, the $r$ parameter needs to be reduced, as we expect reads supporting the reference base in admixed normal tissue. For germline variants, both $r$ and $a$ need to be adjusted as normal cells carry reads supporting both of them. We define the variant allele frequency as follows:

$$VAF_{tum}^{som} = \frac{a}{a+r} \tag{14}$$

By combining formulae 14 and 1 we can conclude that the parameter $r$ needs to be adapted to $r'$ by the following formula:

$$r' = a * (\frac{c_{tum}}{ac_{tum}^{som}} - 1) \tag{15}$$

As we do not expect any alternative supporting reads in normal tissue for somatic variants, formula 15 is sufficient to adjust the number of reads for somatic variants.

#### 2.2 Germline variants

For germline variants, however, where reference and alternative reads are expected in admixed normal tissue, we need to make the distinction between $r_{tum}$ and $r_{norm}$ ($a_{tum}$ and $a_{norm}$), defining the numbers of reads supporting reference/alternative in tumor and in normal tissue, respectively. In order to deconvolve necessary adjustments on these four parameters, we make use of four definitions or formulae, which form a linear equation system:

$$\begin{aligned}
&\textit{(I)} & VAF_{tum}^{germ} &= \frac{a_{tum}+a_{norm}}{a_{tum}+a_{norm}+r_{tum}+r_{norm}} \\
&\textit{(II)} & VAF_{norm}^{germ} &= \frac{a_{norm}}{a_{norm}+r_{norm}} \\
&\textit{(III)} & p &= \frac{\frac{a_{tum}+r_{tum}}{c_{tum}}}{\frac{a_{tum}+r_{tum}}{c_{tum}}+\frac{a_{norm}+r_{norm}}{c_{norm}}} \\
&\textit{(IV)} & r &= r_{tum}+r_{norm}
\end{aligned} \tag{16}$$

The linear equation system has the following solutions:

$$\begin{aligned}
r'_{norm} &= r * \frac{-\frac{p-1}{c_{tum}}}{\frac{(\frac{p}{c_{norm}}-\frac{p-1}{c_{tum}})*VAF_{norm}^{germ}*VAF_{tum}^{germ}}{VAF_{norm}^{germ}-1}+(\frac{p-1}{c_{tum}}-\frac{p}{c_{norm}})*VAF_{tum}^{germ}} \\
r'_{tum} &= r - r'_{norm} \\
a'_{norm} &= r'_{norm} * \frac{-VAF_{norm}^{germ}}{VAF_{norm}^{germ}-1} \\
a'_{tum} &= \frac{-a'_{norm}*(VAF_{tum}^{germ}-1)-VAF_{tum}^{germ}*(r'_{tum}-r'_{norm})}{VAF_{tum}^{germ}-1}
\end{aligned} \tag{17}$$

The formulae in (17) are the required adjustments of the numbers of reference and alternative reads for germline variants. For computation of the genotype likelihood, only $r'_{tum}$ and $a'_{tum}$ are needed. In the implementation in ZygosityPredictor, in order to follow a conservative and safe strategy, the $r'_{tum}$ and $a'_{tum}$ worst reads are selected.
